# An automated model reduction tool to guide the design and analysis of synthetic biological circuits

**DOI:** 10.1101/640276

**Authors:** Ayush Pandey, Richard M. Murray

**Affiliations:** Department of Control and Dynamical Systems, California Institute of Technology, Pasadena, CA

## Abstract

We present an automated model reduction algorithm that uses quasi-steady state approximation to minimize the error between the desired outputs. Additionally, the algorithm minimizes the sensitivity of the error with respect to parameters to ensure robust performance of the reduced model in the presence of parametric uncertainties. We develop the theory for this model reduction algorithm and present the implementation of the algorithm that can be used to perform model reduction of given SBML models. To demonstrate the utility of this algorithm, we consider the design of a synthetic biological circuit to control the population density and composition of a consortium consisting of two different cell strains. We show how the model reduction algorithm can be used to guide the design and analysis of this circuit.

## I. Introduction

Model reduction is an indispensable tool in most engineering applications where mathematical models are used for design and analysis of systems. It is possible to create arbitrarily large and detailed mathematical models of the different processes due to the high complexity of physical systems. However, it can be challenging to use these models for design of system and analysis of the system response. Hence, it is common to model various processes of interest using a “reduced” order model. A reduced order model is a lower dimensional model that has a simple representation, is computationally much faster, and is easier to use for system design compared to any other complex higher order mathematical model. For biological systems, usually, we do not have enough information about the various mechanistic interactions to model these processes effectively. This limitation is manifested in the lack of information about the parameters of a biological model. Therefore, it is even more important to have reduced models to model various biological processes. There are multiple ways and scales at which biological systems are modeled such as using the mass-action kinetics of the chemical reactions [1], the chemical master equation [2], the chemical Langevin equation [3], or methods from statistical mechanics [4]. Mass-action kinetic based modeling uses reaction rate equations where dynamics of various processes are approximated by modeling the concentration of various species making it possible to create models that are low-dimensional and simple compared to other methods. The most widely used model reduction methods utilize the separation of time scales [5] in a model, or use balanced truncation [6–8] that transforms the original model into a lower dimensional system while preserving the input-output behavior. A combined approximate method using singular perturbation theory and balanced truncation is developed in [9]. Other methods such as parameter sensitivity analysis based reduction [10, 11], optimization based methods [12, 13], lumping of parameters [14] of the model, non-dimensionalization [15], and conservation analysis are also widely used. In [16] some of these different methods have been combined together for an improved model reduction method. Generally, the choice of model reduction is dependent on the kind of application and is driven by the questions we want to answer using a reduced model. See [17–19] for a detailed review of some of the well known model reduction methods with applications to biological systems and the current best practices.

Among the methods based on time-scale separation, the Computational Singular Perturbation (CSP) [5] algorithm is one of the most popular numerical methods that identifies the fast and slow time-scales for a model. But, apart from being stiff and laborious to solve, it is not always clear for many applications how fast and slow time-scales can be separated. The Intrinsic Low-Dimensional Manifold (ILDM) [20, 21] is another numerical method based on time-scale separation in which the original model is linearized and then the Jacobian of the resulting linearized system is transformed to identify slow and fast states. But, state transformations are introduced in this method due to which the structure and the physical meaning in the original model are lost. Although balanced truncation based methods can give good error bounds, we face a similar problem since the transformations introduced might lead to the loss of the structure of the original model. Hence, it would be difficult to use such methods to guide the design of biological systems where experimental connections of the model states and parameters are important.

On the other hand, Quasi-steady state approximation (QSSA) [22–25] approach is commonly used in practice to account for experimental connections. The theory driving QSSA based reduction is derived from singular perturbation theory [26] of nonlinear dynamical systems. Our proposed algorithm is motivated by singular perturbation theory, discussed in more detail later. However with QSSA, one needs to be careful when collapsing states into algebraic relations, as it might not always give correct results. A counterexample in [1] elucidates this in detail. The rigorously correct way of performing a QSSA would be to formulate it in singular perturbation theory framework. But, the conditions with singular perturbation are often too strict that it is not possible to formulate the model in this framework for many applications. Hence, it is important to carefully navigate this gap between the rigorously correct singular perturbation and using QSSA directly. Our algorithm in this paper works within this gap and finds reduced order models that minimize the error in desired outputs ensuring robust performance of the reduced models.

In this paper, we present a structured automated model reduction algorithm that systematically finds a reduced model while retaining the structure of the original model. We start by briefly discussing some of the preliminary results followed by problem formulation in Section II before presenting our main results in Section III. We first present the results for systems with linear dynamics and then move on towards the extension of the results for nonlinear dynamics. Subsequently, we give an algorithm that builds on the developed model reduction procedure for nonlinear dynamics. In Section IV, we consider two examples to demonstrate the utility of the proposed algorithm. First, we consider a simple example of a toggle switch circuit and then present the model reduction of a population density and composition control circuit. Our algorithm plays an important role in identifying a reduced model that could be used for design and analysis of this circuit.

## II. Problem Formulation

We denote an eigenvalue of a matrix *P* by *λ*(*P*). The maximum eigenvalue will be denoted by *λ*_max_(*P*). For a statedependent matrix *P*(*x*) we denote the maximum eigenvalue of *P* over all values of *x* by *λ*_max_x__ (*P*). Throughout this paper, we consider the Euclidean 2-norm for vectors. For example, we use the notation ∥*x*∥for the 2-norm of 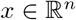 and similarly ∥·∥ represents the induced 2-norm for matrices.

### Definition 1 (*Full model*).

For a state vector 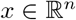, a vector consisting of all model parameters 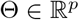 and the vector of output measurements 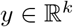, we define the nonlinear dynamic model

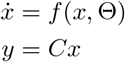

for initial conditions 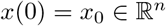.

### Definition 2 (*Reduced model*).

For a full model as defined above, we define a reduced model with a lower dimensional state vector 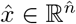 where 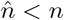 as

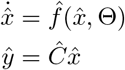

for initial conditions 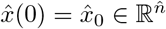 and 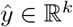.

Note that in the reduced model we have reduced the number of states (from *n* to 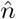) but we have the same number of outputs that depend only on the reduced states. It is clear that in the reduced model some parameters from the full model will be lumped together, hence reducing the number of parameters in the model as well. We will see later how this structure of the reduced model is beneficial for synthetic biological circuit models. Further, we will need the following definitions to formulate the model reduction problem that we solve in this paper.

### Definition 3 (*Sensitivity coefficients*).

For any state, *x_i_* for all *i* = 1,2, …, *n*, the sensitivity coefficient for the state with respect to a parameter *θ_j_* ∈Θ ∀*j* = 1, 2, …, *p* is defined as

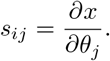

We also define 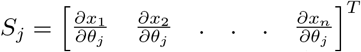, the vector of sensitivity coefficients for all states with respect to the parameter *θ_j_*

### Lemma 1.

*The sensitivity coefficients satisfy a linear differential equation given by,*

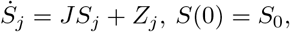

where 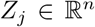 consists of partial derivatives 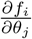 and 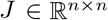 is the Jacobian matrix,

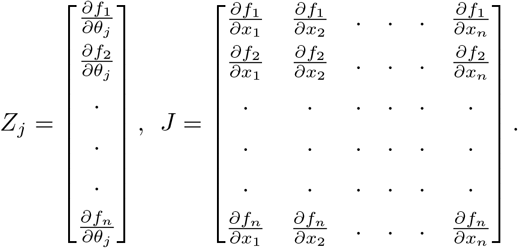

*Proof*. Follows from chain rule of differentiation [27].

Similarly, for the reduced model we can define 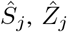 and 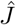 and we have,

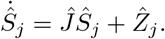

*Remark*. The initial conditions for the sensitivity coefficients

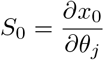

will be equal to zero if the initial conditions *x*_0_ are independent of parameters. If the initial conditions *x*_0_ are dependent on parameters then the corresponding initial condition for the sensitivity coefficients will be equal to 1.

As discussed in the previous section, in our approach we carry over the essential information from the full model to the reduced model by collapsing some of the states’ dynamics into algebraic relations. This process is reminiscent of singular perturbation based model reduction algorithms. In the next theorem, we briefly introduce this topic and refer the interested reader to [26] for more theoretical details and [5] for its applications to model reduction.

### Theorem 1.

*Suppose we have*

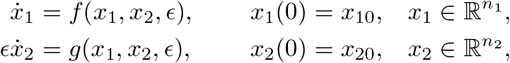

*for* 0 < *ϵ* ≪ 1 and both functions *f* (*x*_1_, *x*_2_, 0), *g*(*x*_1_, *x*_2_, 0) *are well defined. With these conditions, if we are only interested in the slow time-scale dynamics* (*i.e. x*_1_), *we can write the reduced model for this system described by the set of differential equations above. Then,*

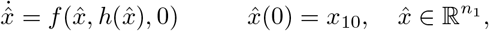

*describe the fast dynamics of the system where x*_2_ = *h*(*x*_1_) *is called the slow manifold. If the eigenvalues of the Jacobian obtained by linearizing* g(*x*_1_, *x*_2_) *with respect to x*_2_ *on the slow manifold have negative real part, then there exist positive constants ϵ* and M such that*

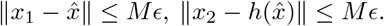

*provided ϵ < ϵ*. Hence, the reduced model converges locally to the full model in the limit.*

*Proof*. See [26].

In the model reduction algorithm that we present in the next section, we use the following definition for the error in reduction and the sensitivity of the error.

### Definition 4.

We define the error in reduction for the output *y_i_* as 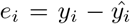 for all *i* = 1, 2, …, *k*. We also define the sensitivity of this error with respect to the model parameter *θ_j_* ∈ Θ as 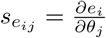 for all *j* = 1,2, …, *p*.

We formulate the desired properties of the reduced models with a view towards how we would like to use these reduced models in design and analysis of synthetic biological circuits. There are two major ways in which reduced models are useful. First, we would like to have simple models that represent the design of a circuit reasonably well so as to use these models to get insights about and improve the design of the various components of the circuit. Secondly, since it is often the case for biological circuits that we can only observe a few states (for e.g. fluorescent outputs), hence, models are often non-identifiable. Reduced models with lower number of states and lumped parameters would decrease the number of non-identifiable parameters in a model. With these applications in mind, we formulate our assumptions accordingly for the model reduction algorithm that we develop.

Given the dynamics of a full model, our objective is to minimize the weighted norm of the error *e* (as in Definition 4) for all time. But, if for a given model we have multiple reduced models that satisfy the desired error tolerance, we would like to have a reduced model that is robust to the uncertainties in the model parameters. We would also like the reduced model to have the least number of states and parameters, to the extent possible. The model reduction algorithm presented in this paper takes into account all of these objectives.

## III. Results

### A. Linear dynamics

Consider the case when the dynamics of the full model are linear of the following form,

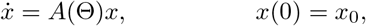

and we want to construct,

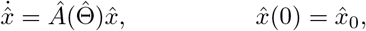

where we assume that *A* and 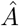 are stable (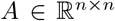 and 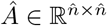) and dependent on the parameters. For outputs, we have *y* = *Cx* and 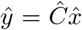. To construct the reduced models, we can choose the states that we would like to collapse and replace their dynamic equation by an algebraic relationship and create a vector of all such collapsed state variables *x_c_*. This procedure to obtain model reduction is similar to the singular perturbation based model reduction in Theorem 1. We can write

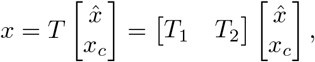

where *T* is a permutation matrix consisting of only zeros and ones with the condition that there can only be one non-zero element in a row and a column. Note that here 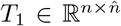 and 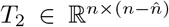. With this structure for *T*, we ensure that the states do not lose their meaning in the reduced model and reduced states are a strict subset of the states of the full model. The model reduction problem then is that of choosing the matrix *T* so that the error in reduction is minimized and the reduced model is robust to parametric uncertainties. Towards that end, we will first consider only the minimization of the two norm of the error for all time. We use the results from [28] for this case. Subsequently, we extend these results to improve the model reduction algorithm so that the performance of the reduced models is robust to uncertainties in parameters.

Define 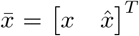, then we can write *e* from Definition 4 as 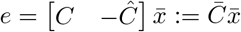. We can write the dynamics of 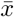,

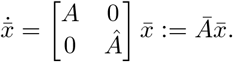

Our objective is to minimize 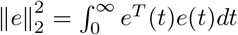 from an initial condition 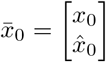.

#### Theorem 2.

*Suppose that there is a matrix P* = *P*^T^ ≻ 0 *that solves the continuous-time Lyapunov equation for the augmented system, that is* 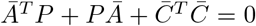, *then the norm of the error in reduction can be upper bounded by the largest eigenvalue of a matrix Q,*

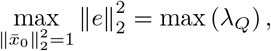

*where*

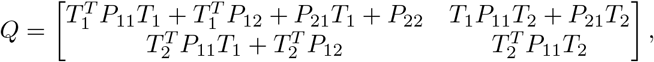

*where we partition *P* as,*

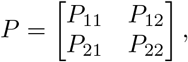

*so that* 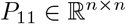 *and* 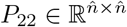.

*Proof*. See [28].

The above theorem gives us an upper bound on the error in model reduction and can be formulated as an optimization problem to solve for the “best” reduced model out of all possible permutation matrices *T*. With the next theorem, we will extend this result to improve the model reduction algorithm so that we are able to choose the reduced models that are least sensitive to uncertainties in parameters. We begin with a lemma that gives us the dynamics of the sensitivity of the error. We use this for the minimization of the sensitivity of error for the linear case.

#### Lemma 2.

*The sensitivity of error (Definition 4) can be described as an output of the linear dynamical system of the augmented sensitivity vector.*

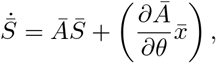

where 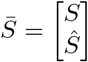, then the sensitivity of the error is given by,

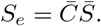

*Proof*. Using the results from lemma 1 for 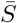,

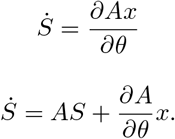

Similarly,

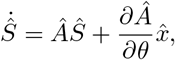

which gives 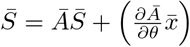 and

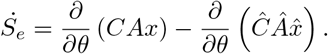

Simplifying, we get the desired result, 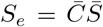 and 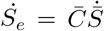.

For the next step of our algorithm, our goal is to minimize 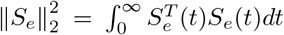. In order to implement this we have to perform the sensitivity analysis of the full model and all possible reduced models for all time to get 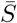 and then use the previous lemma to associate the error sensitivity cost with each reduced model. Clearly, this procedure would be computationally very inefficient. The next theorem gives us a result to simplify this computation by giving a simpler way to get an upper bound on the norm of the sensitivity of the error. We can use the results of this theorem to compute this bound for all possible reduced models and then make a conclusion about the best reduced model along with the results obtained for the minimization of the error.

#### Theorem 3.

*Suppose that there is a matrix P* = *P^T^* ≻ 0 *that solves the continuous time Lyapunov equation* 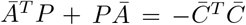 *for the augmented system, then the norm of the sensitivity of the error can be upper bounded as follows,*

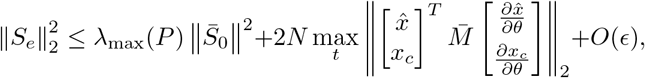

*where *N* is a positive constant, and*

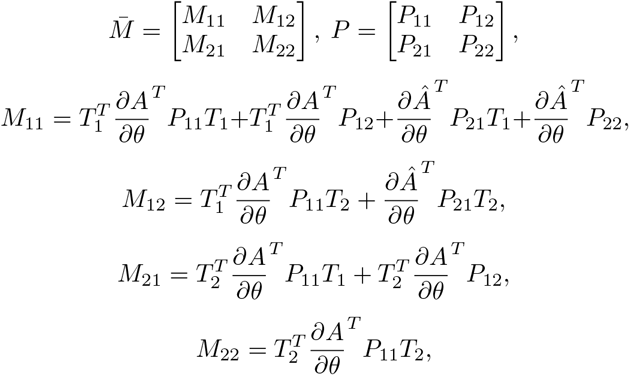

where 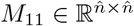 and 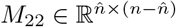.

*Proof.* We prove this theorem in the more general case of nonlinear dynamics in the next section. The proof for the linear case will follow as a special case (shown in Appendix VI-A).

### B. Nonlinear dynamics

For error minimization in the case of nonlinear dynamics, we no longer have the simple analytical results as in Theorem 2. For networked systems with large *n*, i.e. for models with a large number of states the combinatorial procedure of forming possible reduced models and then calculating the error in reduction will be an inefficient algorithm with exponential time-complexity. A greedy algorithm method based on SOSTOOLS [29] to bound the error in reduction for such systems is given in [28]. It gives sub-optimal results and an approximate bound on the error but it is computationally efficient for such systems. However, for our application, we consider minimal ODE models that use Hill functions to approximate the reaction model dynamics in a way that is suitable for experimental design and analysis. With these models, we often do not have a large *n*. Moreover, usually, the states are directly related to the different components of the circuit that we are designing and hence it would not be wise to take a greedy algorithm approach with these models. So, for system dynamics where *n* is not large (< 20), we can computationally calculate the error in model reduction for all possible reduced order models in a brute force way. We discuss this in more detail later when we present the algorithm.

The second part of the model reduction method that we presented for the linear case is the minimization of the sensitivity of the error in model reduction. Using this, we can identify the reduced models that have robust performance in the presence of parametric uncertainties. For the nonlinear dynamics, we have similar results as obtained earlier for the linear case that we now present beginning with the next lemma that shows the sensitivity of error as the output of the augmented sensitivity system.

#### Lemma 3.

*The sensitivity of error as defined in Definition 4 for the nonlinear dynamics can be described as an output of the linear dynamical system of the augmented sensitivity vector,*

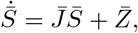

*where* 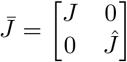 *and* 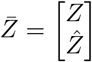. *The error in sensitivity is then given by*,

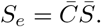

*We can also write the dynamics of the sensitivity of the error* 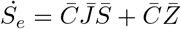.

*Proof*. The proof follows similar to the proof for Lemma 2 where now we have the Jacobians (*J* and 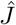) of the system dynamics and the matrices (*Z* and 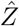) as defined in Section II instead of *A*, 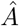 and 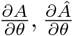.

It is possible to use this lemma to implement an algorithm that solves for the sensitivity of the error by integrating the error sensitivity dynamics equations along with the system dynamics. However, with the next theorem we show a result that gives a method to upper bound the norm of the sensitivity of the error that can be computationally more efficient than the brute force sensitivity analysis of all possible reduced order models.

#### Theorem 4.

*Suppose that for every point* 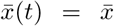 *in the augmented nonlinear system trajectory there is a matrix* 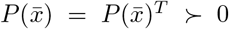 *that solves the continuous-time Lyapunov equation for the augmented Jacobian matrix 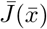, i.e., 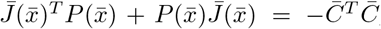, then the norm of the sensitivity of the error for the nonlinear model reduction can be upper bounded by:*

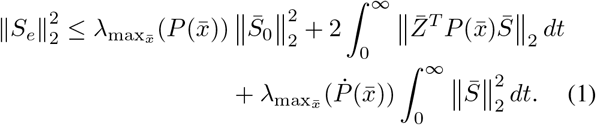

*Proof.* See Appendix A.

### C. Algorithm

In this section, we present a brute force algorithm based on the results presented in the previous section to compute the sensitivity of the error efficiently for all reduced models. Note that the set *R* in the algorithm is the set of all possible combinations of states that we attempt to reduce. This set is created by simply creating combinations of all states but also ensuring that the states corresponding to the outputs are never collapsed. For the purpose of implementation, the permutation matrix *T* in the algorithm consists of indices of states that will be collapsed (as opposed to the 0s and 1s as used above for the theoretical results, where 1 corresponded to the states that were collapsed). The time-complexity of the algorithm is exponential with *n*, the size of the full model, since we search for the best reduced model over all possible combinations of the matrix *T*. However, it is possible to implement the algorithm efficiently when *n* is not large, that is for system dynamics that are not networked dynamics with a large number of states. Particularly, an estimate of the number of states of the full model for which it would be possible to use this algorithm would be *n* < 20. For our application of model reduction of synthetic biological circuit models, this algorithm is suitable, since we often have coarse-grained models that use Hill function modeling [1] to obtain simple enough models that describe the system dynamics effectively. We will demonstrate the utility of the algorithm in the next section with a simple example first and then also for a circuit that implements population density and composition control of two cell strains in a consortium.

#### Algorithm 1: Automated model reduction algorithm

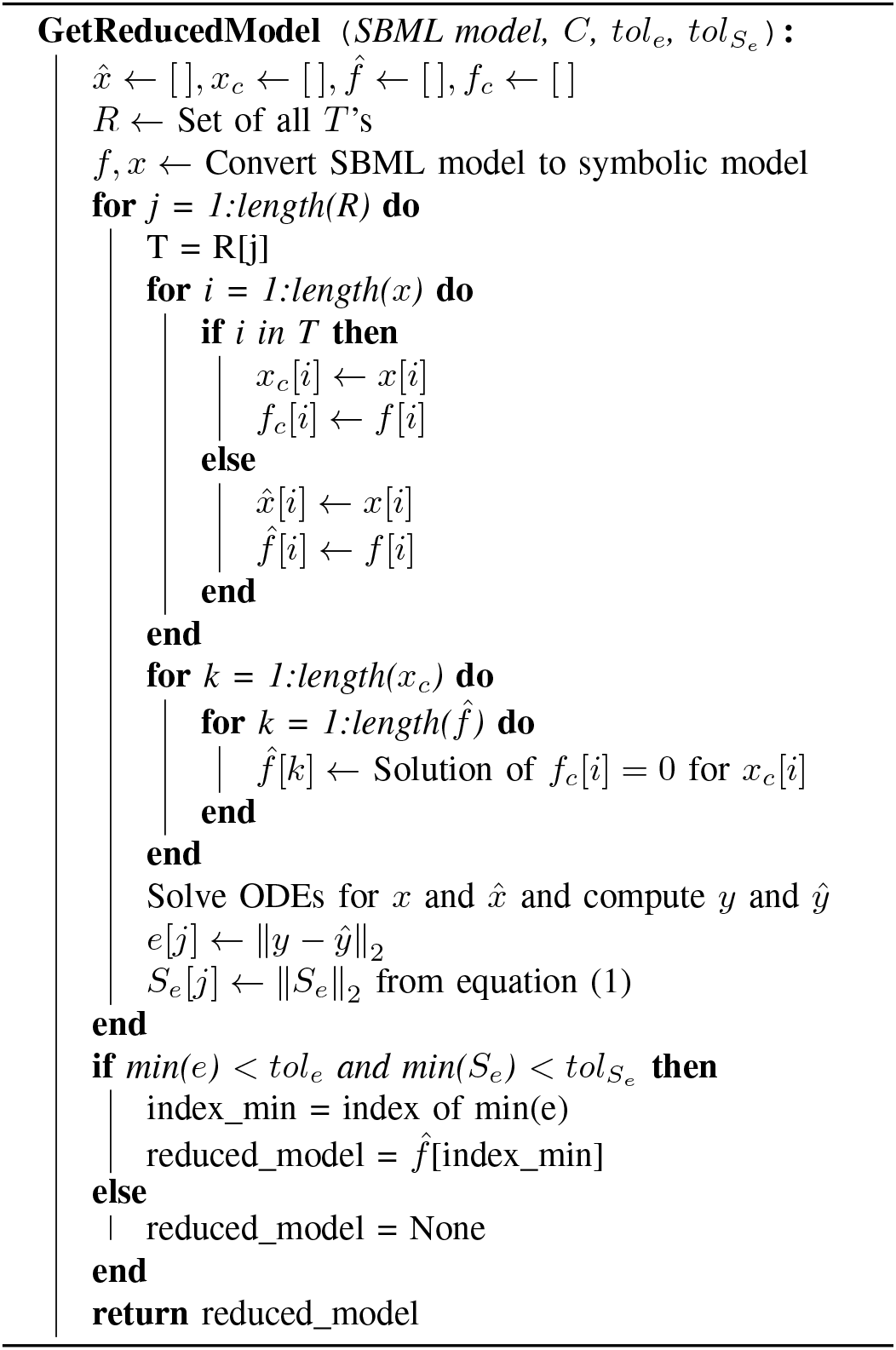

## IV. Examples

We will start by considering a simple example of a toggle switch model.

### A. Toggle Switch

#### 1) Model

The toggle switch is a two species system where both proteins A and B repress each other. We consider the following toggle switch system where proteins TetR and LacI repress each other’s expression. LacI binds with the promoter for protein TetR to prevent the transcription of the TetR gene to form its mRNA and hence repressing the production of protein TetR. Similarly, TetR binds with the promoter of the LacI gene to prevent its transcription to form mRNA, which in turn causes the repression in the expression of LacI protein. This simple two species transcriptional regulation system is shown in Fig. 1. It shows bistability — has two stable equilibrium points and one unstable equilibrium [1]. Here we consider one of the stable equilibrium points and attempt to get a reduced model around this equilibrium. The derivation for the nonlinear ODEs of this system is given in [1]. The final model is

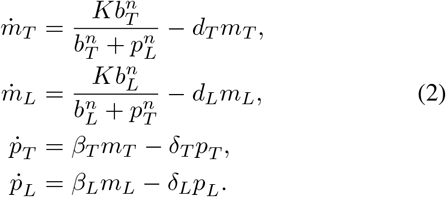

**Fig. 1.**
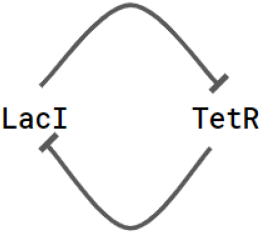
A two species toggle switch where LacI and TetR repress each other’s expression. We can control the strength of each repression using inducers, which effectively control the values of the parameters *b_T_* and *b_L_* in the model given in equation (2)

If we consider the two proteins as output species then

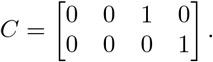

According to the algorithm proposed in the previous section, the possible number of reduced models will be three. The corresponding list *R* will be {(1, 3, 4), (2, 3, 4), (3, 4)} where 1 corresponds to the state *m_T_* and so on for the other three states. For a particular set of parameters where *β_i_, δ_i_* ≪ *b_i_, d_i_*, we can express the model in equation (2) in the singular perturbation theory framework defined earlier in Theorem 1. From singular perturbation theory based reduction, we get the following reduced model (See [1])

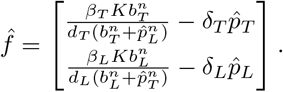

We can use this as a proposed reduced model to validate with the algorithm and the theory we developed in this paper. Particularly, we will see how the conditions when this reduced model is valid imply that the sensitivity of the error in reduction converges to zero. The singular perturbation based reduction is a special case of the model reduction approach using the algorithm presented in the previous section. For this special case, we will see that the error sensitivity with respect to parameters takes a simple form that we can evaluate.

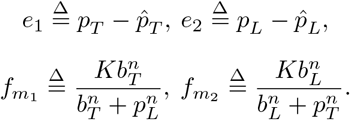

For a parameter *θ_j_* ∈ Θ and *i* = 1, 2, we have,

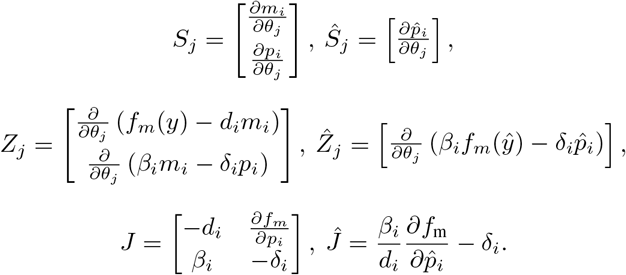

Now, using Lemma 3, we get,

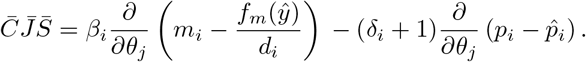

Also from the Lemma 3, for *θ_j_* = *β_i_*, we have 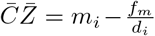 and

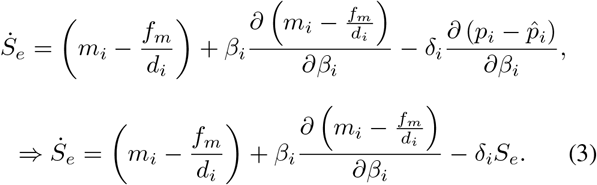

For *θ_j_* = *δ_i_*, we have, 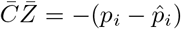,

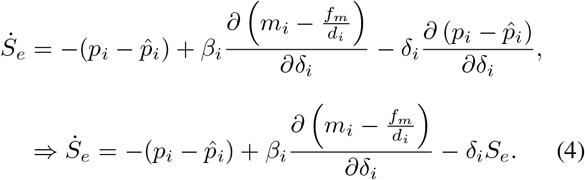

For *θ_j_* = *d_i_*, 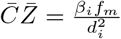,

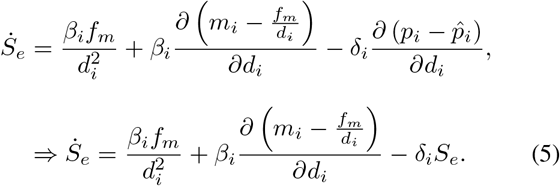

Hence, using equations (3), (4) and (5), the sensitivity of the error in model reduction converges to zero if,

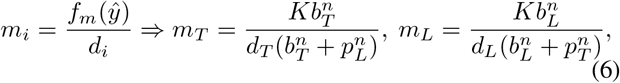

and trivially, the other condition is that the outputs are alike 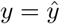.

We run the full model in equation (2) through the automated model reduction algorithm with the desired tolerance levels for the error and the sensitivity of the error. The reduced model returned by our algorithm is given below. It is evident that in this case the reduced model is the same as that given by the singular perturbation theory:

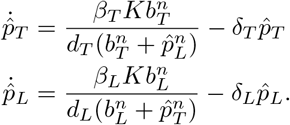

However, if we tune the error margins and desire lower tolerance levels then the algorithm returns a different reduced model which would be one with additional states. It would be difficult to derive such a reduced model using singular perturbation approach. Hence, we can use the automated model reduction algorithm to control the tradeoff between the number of states and error in model reduction along with the sensitivity of the error to find reduced models that would be otherwise elusive to find using other existing methods. We will see with the next example that minimizing the error in reduction is not enough as it returns a set of reduced models that all have similar error bounds. We will use the automated model reduction algorithm to find out the reduced model that has the best robust performance in presence of parametric uncertainties to choose one final reduced model out of the multiple possibilities we might have when minimizing only the error.

### B. Population and composition control circuit

In this example, we consider a synthetic circuit designed to control a consortium of two different cell strains. Two coupled feedback controllers are designed using various components that control the total population density of the consortium to a desired value and to control the fraction of the two cell types in the consortium. In [30], the theory behind the design of these controllers has been discussed and [31] discusses the details of the implementation of this circuit. In this circuit, each cell in the consortium promotes the production of a toxin that kills itself. At the same time, each cell also signals the production of anti-toxin in the cells of the other type using signaling molecules that can transport over from one cell to other. The production of anti-toxin rescues the cell from killing itself by binding the toxin in that cell. Amount of two different inducers can be used to control the expression of these signaling molecules. There are two different fluorescent output readouts corresponding to each cell type. Hence, this system has two outputs (*L*_1_ and *L*_2_ in the model).

#### 1) Mathematical model

The model dynamics can be described by the ODE model given in equation (7). The description of the model species and the model parameters are given in Appendix B. For more details on the parameter values and description, the reader is referred to [31].

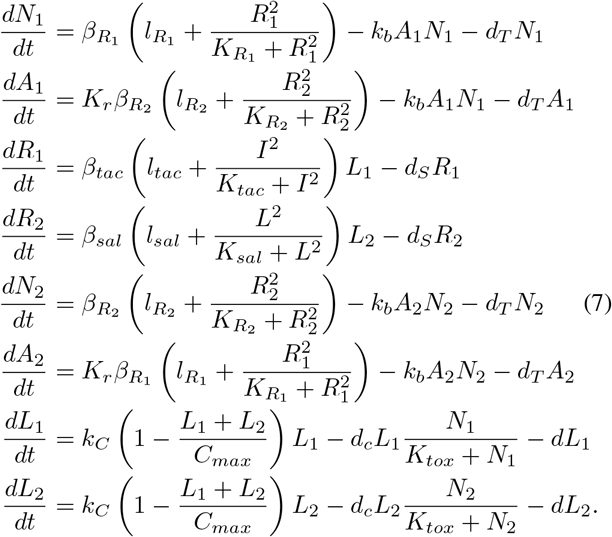

#### 2) Model reduction

For the model of this system, using singular perturbation theory to reduce the model is not possible as it is not clear how this model can be expressed in the singular perturbation framework as defined in Theorem 1. We will show how the automated model reduction developed earlier can be used to reduce the model to get a reduced model that has 4 states, down from a total of 8 in the full model. As discussed earlier, this model reduction could be beneficial in both the design guide for the circuit and the analysis using the model. With the four state reduced model, we can make easier and direct connections with the tunable parameters in the system since there are only two inputs and two outputs in the system. Using simulations, we can study the response of the system for different parameters and make connections with the experimental setup to guide the design of the experiment. Moreover, the control functions implemented in this circuit can be conceptually achieved in multiple ways. For instance, in the circuit described above, both cells kill self and rescue the other. There are other possible configurations that could implement similar control functions in principle (repressing growth/promoting death of the other cell etc.), we need to scan over all such possibilities to determine which one could be implemented experimentally to give the desired results. Using reduced models for such purpose is important as working with complicated models can be disconnected from the control functions whereas with a reduced model this is not the case as much. The advantages of using a reduced model for parameter identification are clear, since, we only have two measurements and 24 parameters in the full model. However, as we will see in the final reduced model there are only 4 states and 13 parameters. Hence, the number of non-identifiable parameters will be much lesser, giving better estimation of parameters.

Using the automated model reduction algorithm for this model, we get four different reduced order model (all with 4 states) that perform similar on the normed error metric. The results are presented in Fig. 2. Running the algorithm with both the minimization of sensitivity of error metric and the minimization of the normed error, we can find out the following reduced model that has robust performance in total population density control.

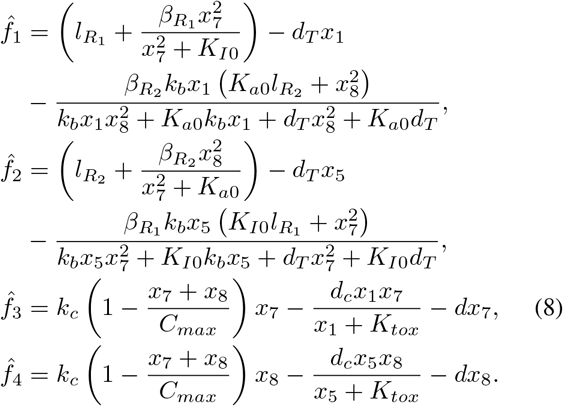

**Fig. 2.**
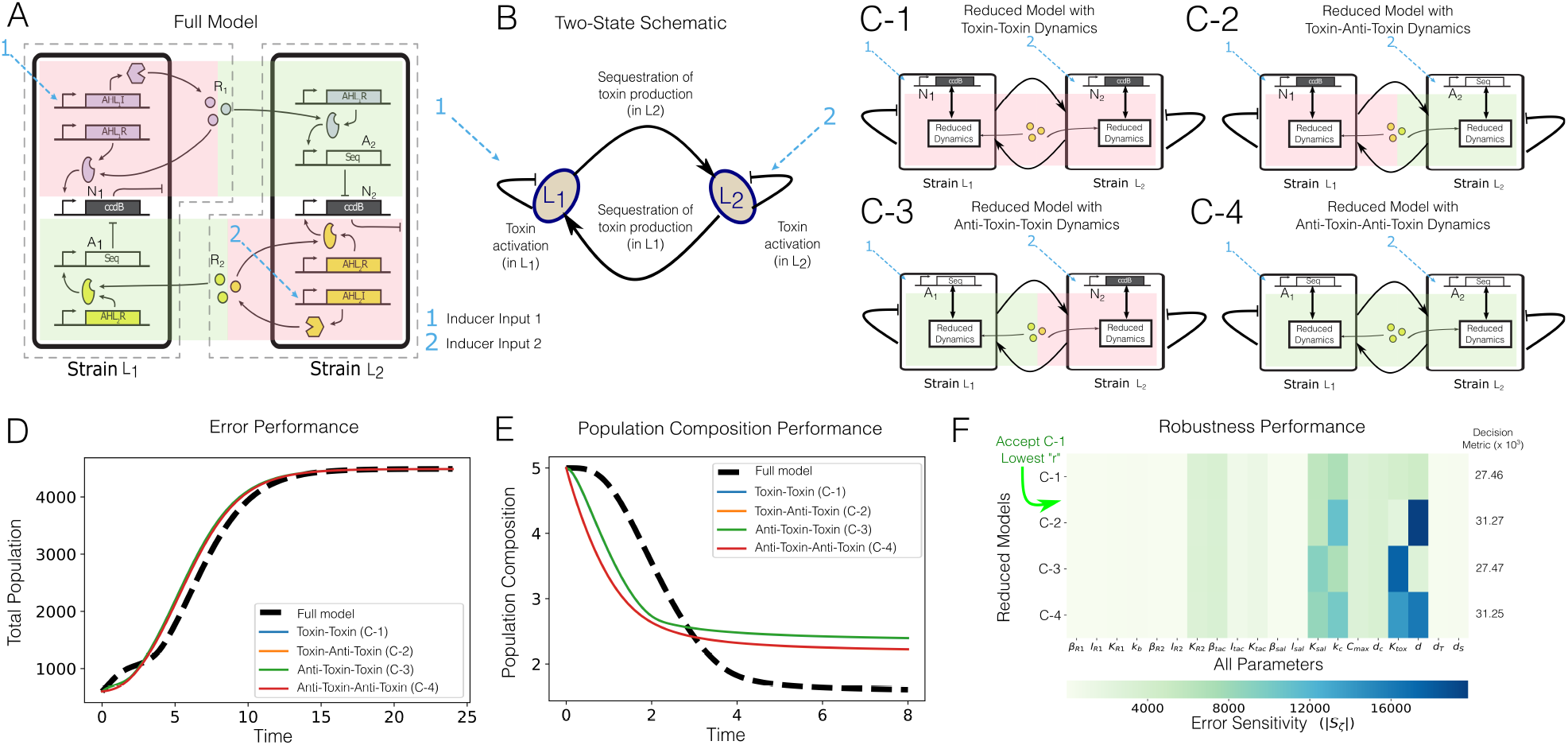
**(A)**: Two-member population and composition control circuit schematic. The two inducer inputs (labeled 1 and 2 in cyan) activate the ccdB expression in *L*_1_ and *L*_2_ cell strains respectively. AHL signals diffuse out of each cell type to signal the expression of anti-toxin (ccdA) in the other cell type. The anti-toxin sequesters away the toxin protein to rescue the cell population. **(B)**: A simple schematic demonstrating the proliferation and death of each cell type under input signals. (**C-1 – C-4**): Reduced models obtained using our automated model reduction approach. Each reduced model has four states. Two of these states are the output signals which are never reduced in our method. The other two states in each reduced model are labeled in the Figure. **(D)**: Total population (*L*_1_ + *L*_2_) obtained on simulating each reduced model and the full model. As we can see, the error performance for all of these four reduced models is satisfactory. **(E)**: Population composition (*L*_1_/*L*_2_) for the reduced models and the full model. Composition control is a feature of this circuit that the reduced models demonstrate as well, although the dynamics of population composition are not fully captured. **(F)**: Robustness analysis for each reduced model gives us a guide to choose a particular reduced model given the parameters in the full model. The heatmap shows the norm of the sensitivity of error in model reduction ∥*S_ζ_*∥ with respect to the model parameters. The decision metric *r* = *w*_1_∥*ζ*∥ + *w*_2_*d_R_* is then used to choose the reduced model given in (C-1) since it has the least *r*

Here 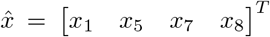 and the lumped parameters are derived in Appendix C.

## V. Conclusion

We proposed an algorithm based on quasi-steady state approximation model reduction that collapses the dynamics of a subset of states of the full model to algebraic relationships. Motivated by applications to reduce synthetic biological circuit models, we developed a structured model reduction algorithm that minimizes the error and its sensitivity between the full model and the reduced model. The algorithm returns a reduced model that not only has similar output responses as the full model but also is robust to parametric uncertainties. Towards that end, we extend the results of [28] by including sensitivity of error in the objective cost to search for reduced models. We show using an example of a toggle switch circuit how this additional constraint relates to the singular perturbation theory based model reduction. Moreover, using an example of a circuit designed to control the population and composition of a consortium consisting of two cell strains we show that error minimization as the only constraint does not give satisfactory results in finding a reduced order model that best represents the full model. For this example, we show the application of our algorithm that minimizes the sensitivity of the error along with the error itself to return a final reduced model that has robust performance to parametric uncertainties. We also give theoretical results that bound the sensitivity of the error both for autonomous linear and nonlinear dynamics with linear output relationships. A direct extension of this work would be to derive results for more general system dynamics with inputs and nonlinear output relationships. It would also be interesting to look for ways to improve the computational efficiency of our algorithm, both by improving the theoretical results and by improving the algorithmic implementation.

## VI. Acknowledgements

We would like to thank Chelsea Hu, Reed McCardell, and Shailja for insightful discussions. We would also like to thank Samuel Clamons for helping with sensitivity analysis computations. The author A.P. is supported by the Defense Advanced Research Projects Agency (Agreement HR0011-17-2-0008). The content of the information does not necessarily reflect the position or the policy of the Government, and no official endorsement should be inferred.

## Appendix

### A. Proof of Theorem 4

*Proof*. At the point 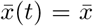 in the augmented nonlinear system trajectory, write the sensitivity system equations [27] for a parameter *θ_i_* ∈ *θ*:

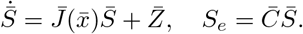

For the norm of the sensitivity of the error, we can write

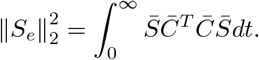

For every 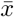, given that there exists a matrix 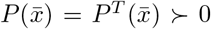 such that 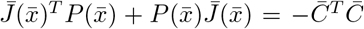, consider a function 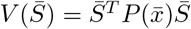. Differentiating this function with respect to time, we have that

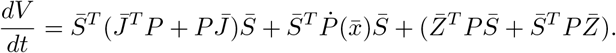

For simplicity of exposition, we denote 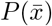 as *P* noting that this is a state-dependent Lyapunov matrix, similarly 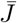 is a state-dependent Jacobian matrix of the augmented nonlinear system evaluated at 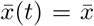. Integrating the expression above from 0 to ∞ and then substituting the expression for 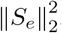, we get

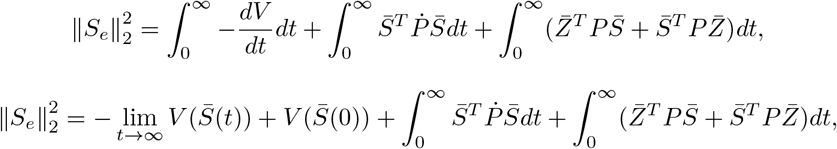

Since, *P* is a positive semidefinite matrix, *V* will be a non-negative function [26]. Using this fact and denoting 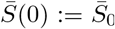, we have the inequality

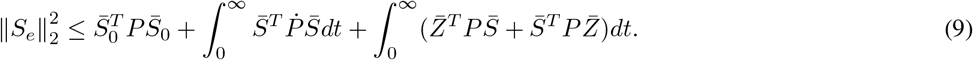

For the first part of this equation, we can write

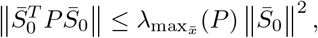

where we compute the maximum eigenvalue of *P* over all points 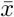. Note that if the initial conditions are independent of all model parameters then 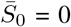. Similarly, for the second term we get,

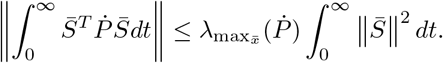

Evaluating the last part in equation (9) proves the theorem, combined with the above results:

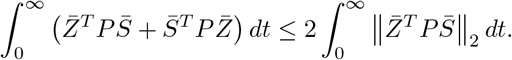

For the special case of linear system dynamics, we can further simplify the bound above since 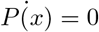 since the Lyapunov matrix is the same at all points in the trajectory and we can asymptotically bound the integral using the stability properties of the system. We have that

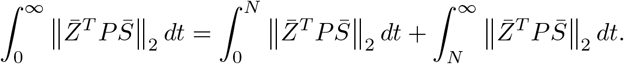

We can show that the tail-end integral is bounded as *O*(*ϵ*) by writing out the sensitivity 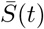 explicitly and using the asymptotic stability of the augmented system. Refer to the Appendix in [32] for the sensitivity equation and its use in bounding ∥*S*∥.

*Remark.* For computational purposes, we may modify the bound above as

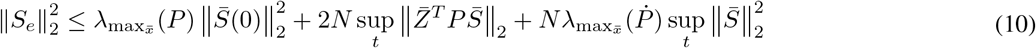

where *N* > 0 is a time at which the system solution is arbitrarily close to the equilibrium point, that is, 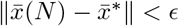 for some *ϵ* > 0 and 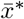 is the equilibrium point for the augmented nonlinear dynamical system. Since we compute 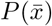 at every time step, we can also compute 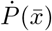 numerically. Further, a direct computation for *S_e_* is also possible but the bounds that we give may be used as an interpret-able metric for system performance analysis.

**TABLE I.** Model parameters

| S.no. | Parameters | Description | Unit | Guess |
| --- | --- | --- | --- | --- |
| 1 | $\beta_{R_1}$ | Max transcription rate of Lux inducible promoter | con./hr | 6 |
| 2 | $l_{R_1}$ | Leak constant of Lux inducible promoter | N/A | 2e-3 |
| 3 | $K_{R_1}$ | Activation constant of Lux inducible promoter | con. | 430 |
| 4 | $k_b$ | Binding rate between toxin and anti-toxin | 1/con.hr | 30 |
| 5 | $\beta_{R_2}$ | Max transcription rate of Cin inducible promoter | con./hr | 6 |
| 6 | $l_{R_2}$ | Leak constant of Cin inducible promoter | N/A | 2e-3 |
| 7 | $K_{R_2}$ | Activation constant of Cin inducible promoter | con. | 30 |
| 8 | $\beta_{tac}$ | Max transcription rate of inducible promoter $P_{tac}$ | con./hr | 19.8e-3 |
| 9 | $l_{tac}$ | Leak constant of inducible promoter $P_{tac}$ | N/A | 1.5e-3 |
| 10 | $K_{tac}$ | Activation constant of inducible promoter $P_{tac}$ | con. | 1.4e5 |
| 11 | $\beta_{sal}$ | Max transcription rate of inducible promoter $P_{sal}$ | con./hr | 14.4e-3 |
| 12 | $l_{sal}$ | Leak constant of inducible promoter $P_{sal}$ | N/A | 2.1e-4 |
| 13 | $K_{sal}$ | Activation constant of inducible promoter $P_{sal}$ | con. | 4.3e4 |
| 14 | $k_C$ | Cell Division rate | 1/hr | 0.6 |
| 15 | $C_{max}$ | Population cap | conc. | 5500 |
| 16 | $d_c$ | Cell death rate | 1/conc.hr | 0.8 |
| 17 | $R_2$ | Max induced Cin concentration | con. | 100 |
| 18 | $I$ | Max induced IPTG concentration | con. | 1e6 |
| 19 | $R_1$ | Max induced Lux concentration | con. | 1e3 |
| 20 | $Sal$ | Max induced Sal concentration | con. | 1e5 |
| 21 | $K_{tox}$ | Repression coefficient of toxin to proliferation | con. | 5 |
| 22 | $d_S$ | Degradation constant of AHLs | 1/hr | 0.1 |
| 23 | $d$ | Basal degradation of each cell | 1/hr | 0.01 |
| 24 | $d_T$ | Basal degradation of toxins and antitoxins | 1/hr | 0.01 |
| 25 | $K_r$ | Ribosome scaling factor | N/A | 5 |

**TABLE II.** Model species

| Species | Description |
| --- | --- |
| $N_1$ | Average toxin (ccdB) con. in $L_1$ population |
| $A_1$ | Average anti-toxin (ccdA) con. in $L_1$ population |
| $R_1$ | Signal 1 ($R_1$), Lux con. in consortia |
| $R_2$ | Signal 2 ($R_2$), Cin con. in consortia |
| $N_2$ | Average toxin (ccdB) con. in $L_2$ population |
| $A_2$ | Average anti-toxin (ccdA) con. in $L_2$ population |
| $L_1$ | Cell type 1 ($L_1$) population count |
| $L_2$ | Cell type 2 ($L_2$) population count |

### C. Model reduction of population control circuit

When the dynamics for *x*_2_, *x*_3_, *x*_4_, *x*_6_ are collapsed into algebraic relationships, i.e. 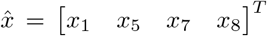, we get the following reduced order model,

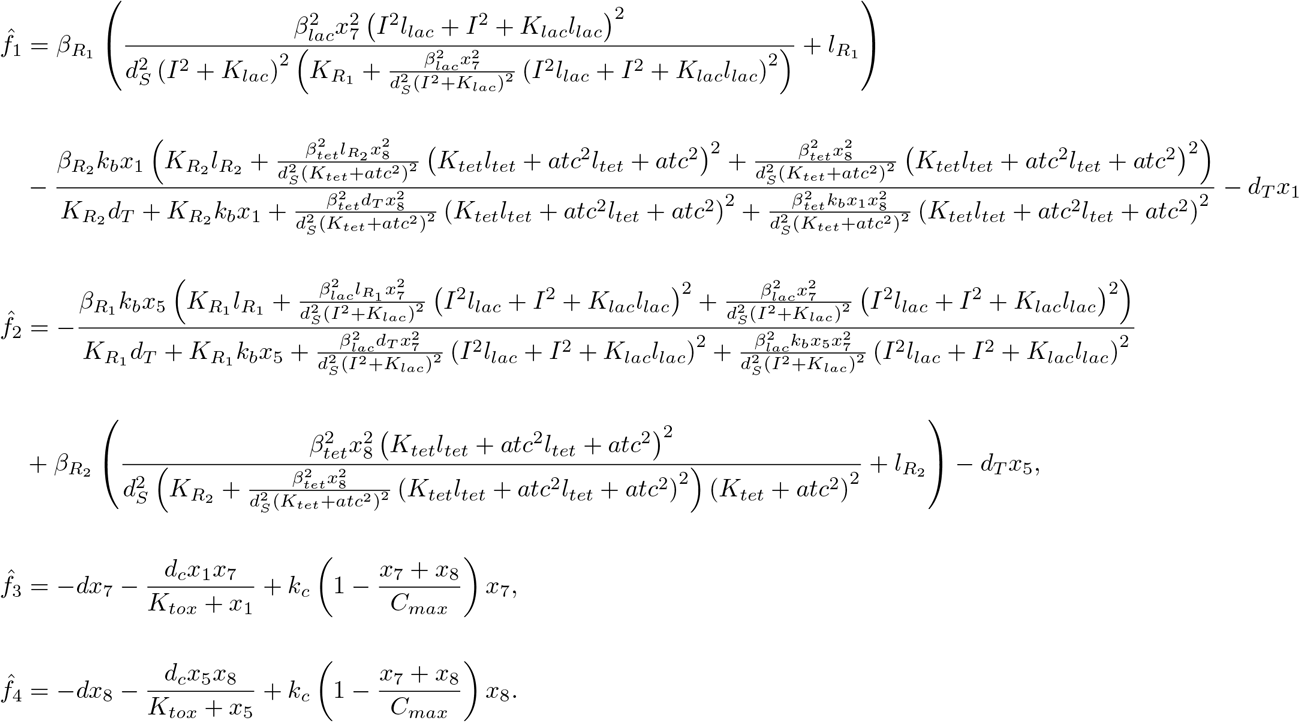

Lumping parameters together, we get

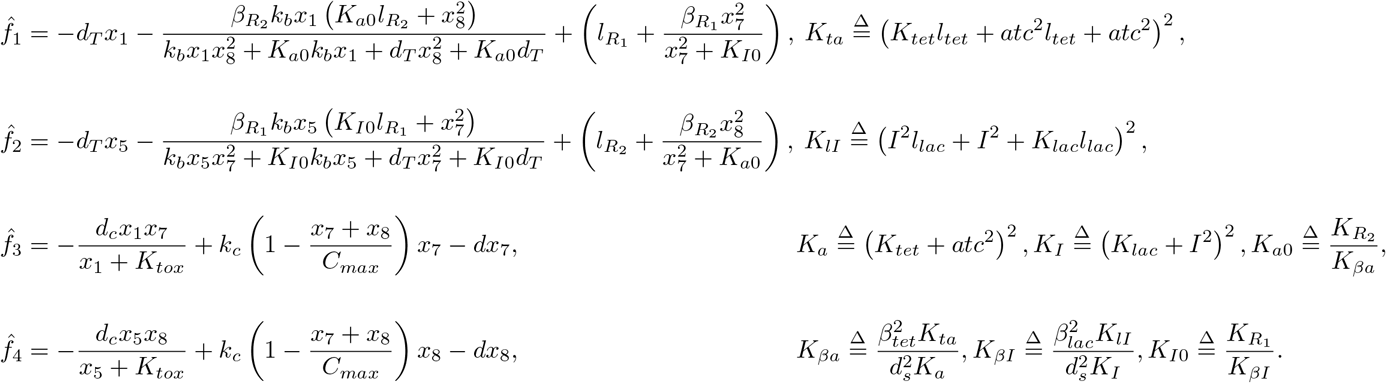

As we can see we have reduced the total number of parameters in the reduced model from a total of 24 in the full model to 13 in the reduced model. Other reduced models with four states are obtained by collapsing the dynamics for (*x*_2_, *x*_3_, *x*_4_, *x*_5_), or (*x*_1_, *x*_3_, *x*_4_, *x*_5_), or (*x*_1_, *x*_3_, *x*_4_, *x*_6_). All four of these reduced models have four states and minimize the error metric. In Fig. 3, we show the sensitivity analysis heatmap for the two outputs of the reduced model derived above with respect to all model parameters. Similar heatmap plots can be obtained for all of the other reduced models, given in Fig. 4. Using this we then conclude that the reduced model in equation 8 has the lowest sensitivity out of all the others and hence has robust performance in the presence of parametric uncertainties.

**Fig. 3.**
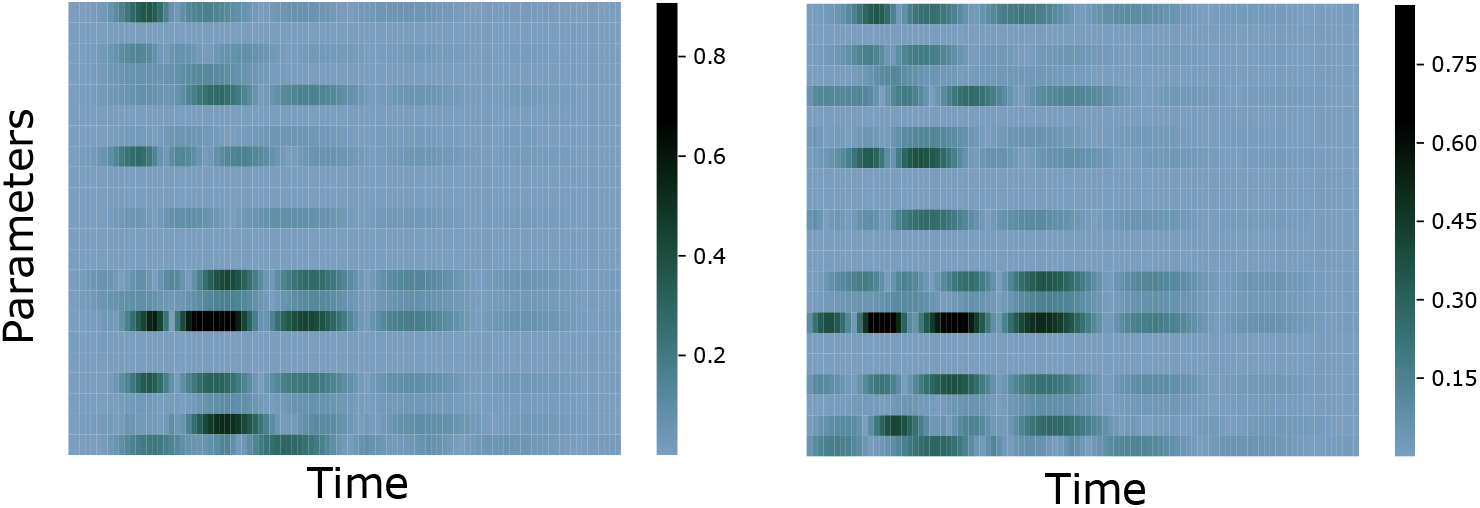
Normalized sensitivity coefficients of the two outputs, *y*_1_ (left) and *y*_2_ (right) corresponding to *x*_7_ and *x*_8_, i.e. the population of each cell type, with respect to the model parameters for the reduced model shown above with 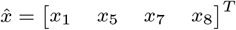.

**Fig. 4.**
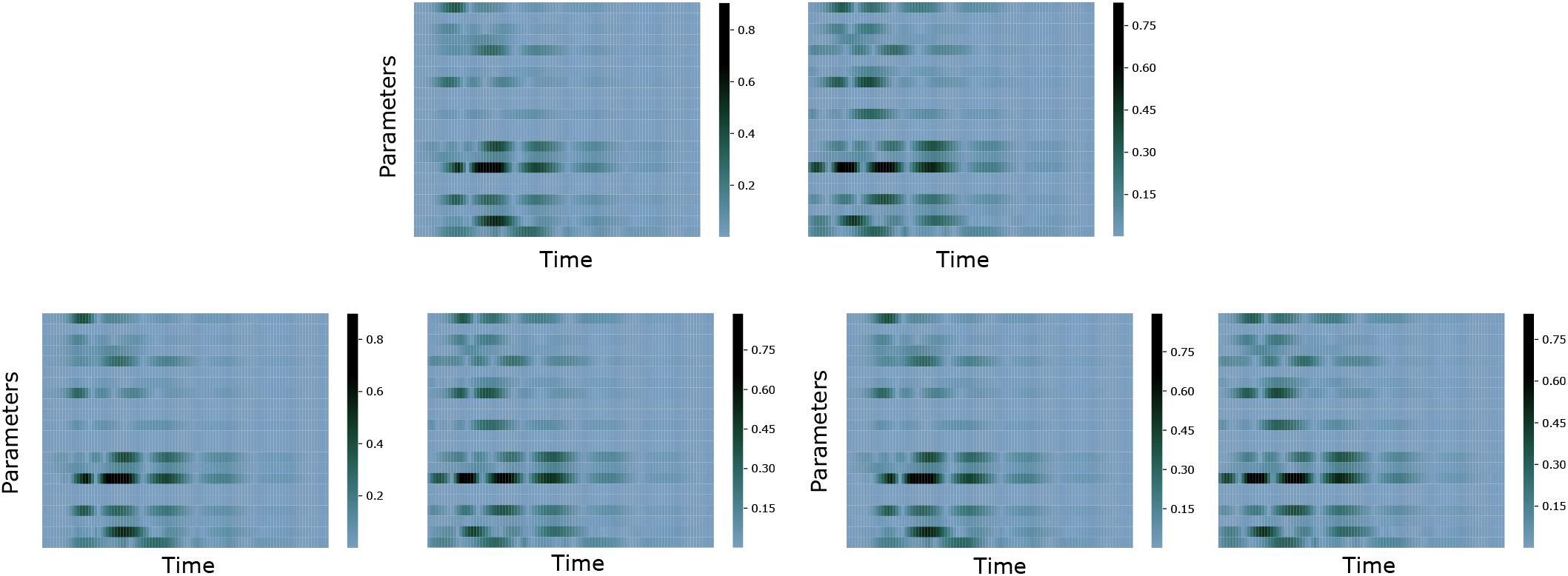
Normalized sensitivity of the two outputs, *y*_1_ and *y*_2_ corresponding to *x*_7_ and *x*_8_, i.e. the population of each cell type, with respect to the model parameters for different reduced models. (top) We have the reduced model with 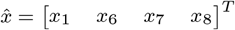. (bottom left) We have the reduced model with with 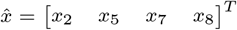 and (bottom right) we have 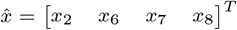. Note that the left plot is the sensitivity analysis for the output *y*_1_ and the right is for the second output *y*_2_ in each case.

